# The effect of spherical projection on spin tests for brain maps

**DOI:** 10.1101/2024.12.15.628553

**Authors:** Vincent Bazinet, Zhen-Qi Liu, Bratislav Misic

## Abstract

Statistical comparison between brain maps is a standard procedure in neuroimaging. Numerous inferential methods have been developed to account for the effect of spatial autocorrelation when evaluating map-to-map similarity. A popular method to generate surrogate maps with preserved spatial autocorrelation is the spin test. Here we show that a key component of the procedure — projecting brain maps to a spherical surface — distorts distance relationships between vertices. These distortions result in surrogate maps that imperfectly preserve spatial autocorrelation, yielding inflated false positive rates. We then confirm that targeted removal of individual spins with high distortion reduces false positive rates. Collectively, this work highlights the importance of accurately representing and manipulating cortical geometry when generating surrogate maps for use in map-to-map comparisons.

## INTRODUCTION

Modern neuroimaging often necessitates evaluating the similarity between brain maps. Technological and data sharing advances have resulted in the proliferation of whole-brain maps of multiple structural and functional features [1], including gene expression [2–4], neurotransmitter receptors [5–8], synapse types [9], cell types [10, 11] and morphology [12], laminar differentiation [13, 14], myelination [15–18], metabolism [19, 20], and intrinsic dynamics [21–26]. Against this backdrop, scientific discovery and inference typically involves a simple yet fundamental step: computing spatial correlations between pairs of brain maps. These comparisons are used to address two broad classes of questions. The first is to contextualize brain maps; for instance, asking whether an experimentally generated map (e.g. a cortical thickness contrast between patients and controls) is enriched for a particular micro-architectural feature (e.g. neurotransmitter receptors) [8]. The second is to identify ontological links between levels of description in the brain; for instance, asking whether a micro-scale feature (e.g. intracortical myelin) is related to a macro-scale feature (e.g. functional hierarchy) [18].

Importantly, brain maps are spatially auto-correlated: areas that are spatially close to one another tend to be more similar than areas that are spatially distant from one another. A consequence of this is that observations are not independent from one another, violating the assumptions of both conventional parametric or non-parametric tests of significance [27, 28]. Specifically, spatial autocorrelation inherent in brain maps has been shown to yield spurious correlations and inflated *p*-values, leading to increased false positive rates [29, 30]. In other words, the background influence of spatial autocorrelation in brain maps can lead to a spurious interpretation that two brain features are related to each other [31]. As a result, multiple methods have been developed to generate surrogate brain maps that attempt to preserve spatial autocorrelation in empirical maps and build null distributions for map-to-map correlation coefficients [16, 28, 32] (for a review, see [31]).

Perhaps the most widely used method to generate surrogate maps with preserved spatial autocorrelation is a spatial permutation procedure that involves rotating spherical projections of brain maps [27, 33]. Specifically, feature values represented on a brain surface are projected to a sphere, subjected to random angular rotation, and then projected back onto the brain surface. Theoretically, this procedure should yield a new brain map in which the original feature values and their spatial autocorrelation are preserved, but their relationship to the underlying brain areas has been randomized. Since its inception, this intuitive and simple procedure — colloquially referred to as the “spin test” — has been readily adopted by the neuroimaging community, leading to numerous variants and implementations [27, 29, 34–3]

Despite its popularity and conceptual appeal, multiple benchmarking studies have shown that in controlled simulated settings, the spin test does not perfectly control for false positive [29, 30]. Here, we explore why this is the case. We first illustrate how the projection of brain surfaces to the sphere leads to deviations from the empirical spatial autocorrelation. We also show that these errors directly result in greater false positive rates. Finally, we show how targeted removal of high-error surrogate maps can be used to reduce the false positive rates.

## RESULTS

### Spherical projections inflate false positive rates

The main goal behind rotating the brain is to generate surrogate brain maps that preserve the spatial structure of the original data. We start out by developing intuition about why projecting surface maps to a sphere distorts distance relationships between vertices. Fig. 1 (left) shows two sets of equally distant vertices on a brain surface, shown in red and blue. Projecting a brain surface causes distortions [38], approximately preserving some distance relationships (red) but disrupting others (blue). A rotation on the sphere is an isometry, meaning that it is a distance-preserving transformation. As a result, rotations of the spherical representation perfectly preserve distance relationships among the vertices (Fig. 1, middle). Projecting the spherical representation back to a brain surface then erroneously preserves distortions and introduces additional ones (Fig. 1, right). In other words, the entire procedure does not necessarily preserve distance relationships in the brain.

**Figure 1.**
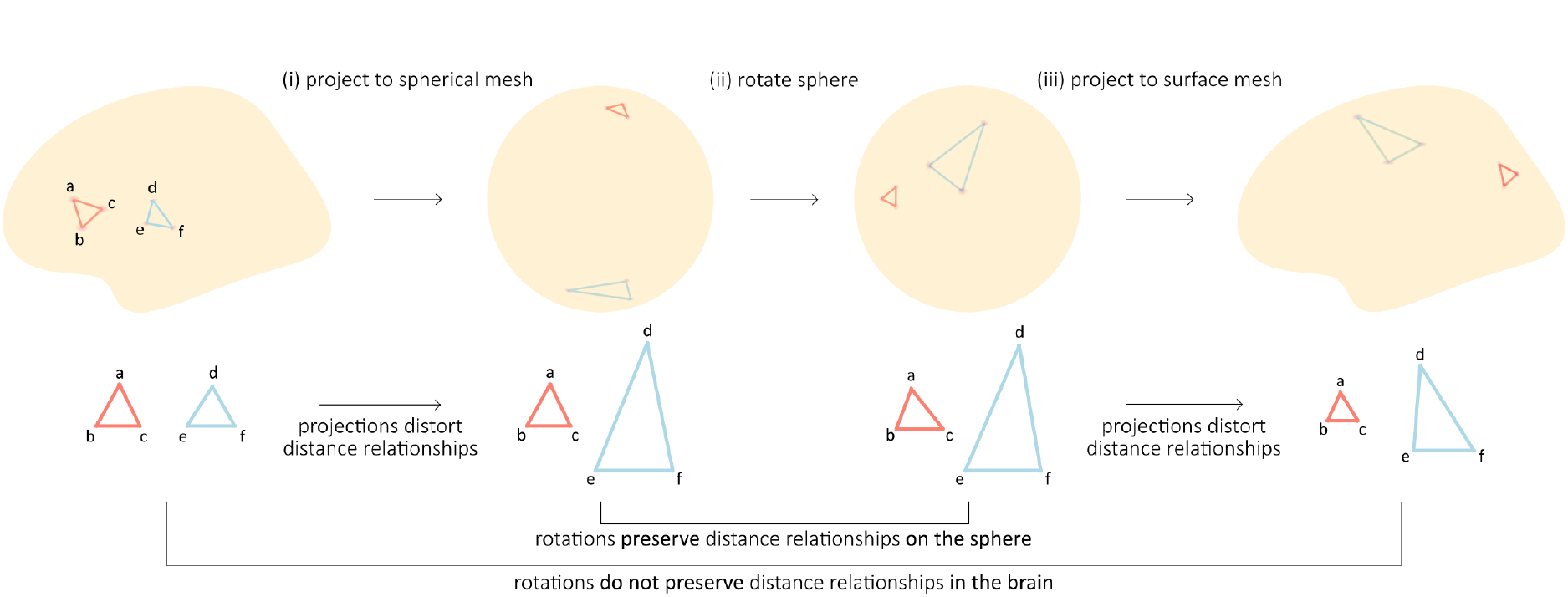
Spherical projections distort distance relationships. The spin procedure consists of three main steps: (i) projecting a brain map to a spherical mesh, (ii) rotating the sphere and (iii) projecting the brain map back to the brain surface. The projection to a spherical mesh distorts distance relationships between vertices of the cortical surface mesh. For instance, the vertices a, b and c (red triangle) as well as d, e and f (blue triangle) are equally distant from one another on the cortical mesh but following the transformation, some distance relationships are not preserved. Namely, vertices d, e and f are significantly more distant from one another, with vertex d farther away from vertices e and f. The rotation of the sphere, being an isometry, preserves distance relationships on the sphere. The projection from the sphere back to the brain further distort distance relationships. For instance, vertices a, b and c are closer to one another, while vertex d is now closer to vertex e than vertex f. Overall, the spin procedure does not preserve distance relationships in the brain.

Fig. 2 demonstrates this point: that rotations are isometries of a sphere but they are not isometries of the brain. To quantify the statistical performance of the spin test, we use ground-truth simulations in which we compute false positive rates (FPRs) when correlating pairs of spatially autocorrelated random maps [28, 29] (see *Methods)*. Briefly, we first generate random maps that are spatially autocorrelated on the sphere using a spatial random field model. By tuning the length parameter of this model, we control the spatial autocorrelation of the generated maps. We generate 1 000 maps with a length parameter ranging from 1mm to 50mm. Then, for each random map, we compute the Pearson correlation between itself and the remaining 999 maps, and evaluate the significance of this Pearson correlation with a spin test, obtaining a total of 999 *p*-values for each random map. Setting the significance threshold to 0.05, we quantify the false positive rate of the statistical test for each random map.

**Figure 2.**
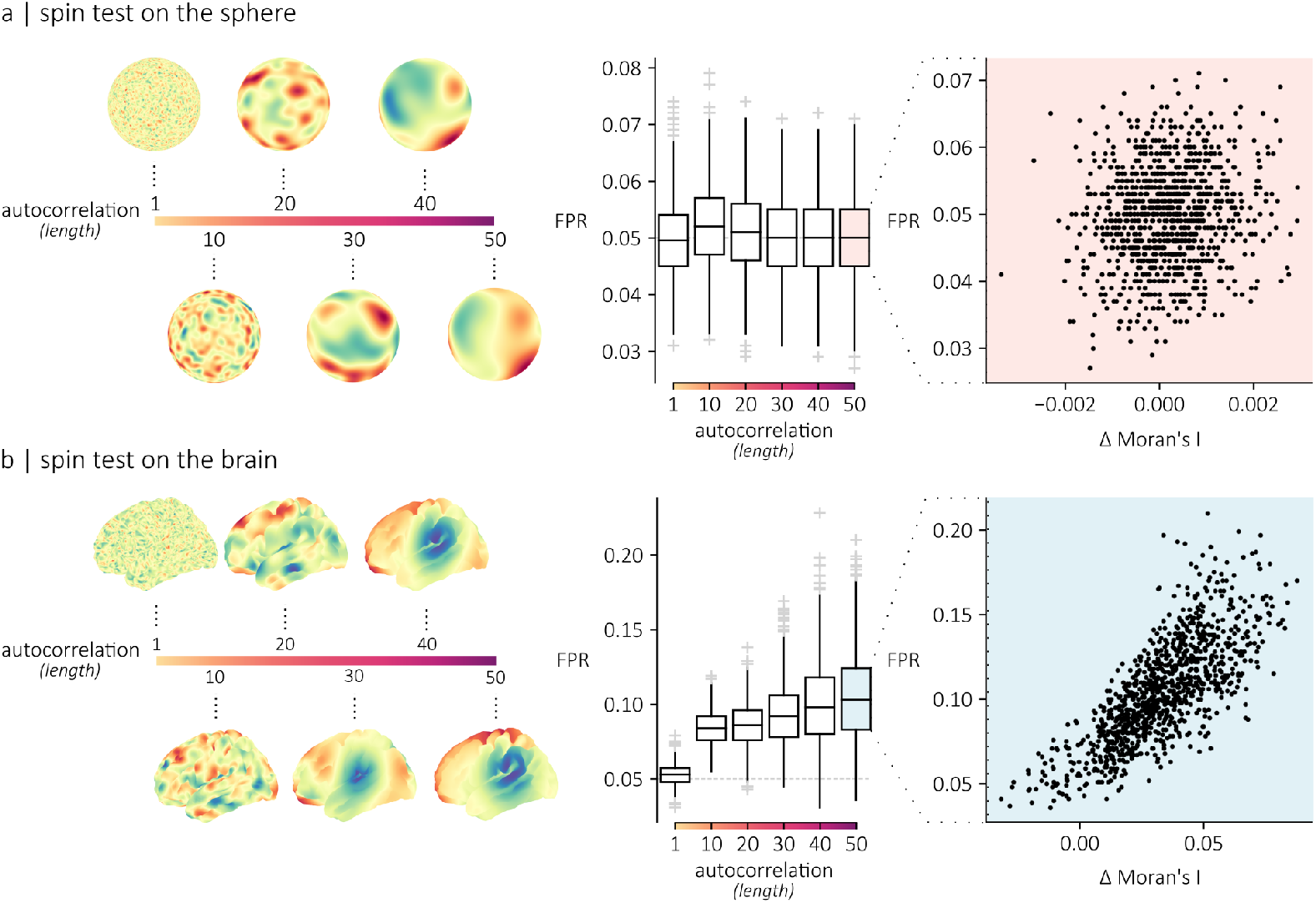
Spherical projections inflate false positive rates. The statistical performance of the spin test was evaluated using spatially autocorrelated maps generated by a Gaussian covariance model with a length parameter defining the spatial extent of the process. A sample of 1 000 maps were generated for six different levels of spatial autocorrelation (lengths) on either a spherical surface (a) or on a cortical surface (b). For each level of spatial autocorrelation, individual maps were correlated with the remaining 999 maps. The significance of each correlation coefficient was evaluated using the spin test. The false positive rate (FPR) was then quantified for each map individually. The boxplots show the distribution of false positive rate for each map, across lengths. The scatterplot shows the relationship between the false positive rate of a map and its average deviation in Moran’s I relative to its rotated versions (empirical − rotated).

Fig. 2a shows that, on average, the false positive rate for random maps generated on a sphere is 0.05 regardless of the spatial autocorrelation, confirming that the randomization (and thus, the significance test) works as expected. By evaluating the average deviation in Moran’s I between each map and their rotated versions, we also confirm that, on average, the spatial autocorrelation of the original maps is preserved following rotation. Fig. 2b shows the same experiment, but with random maps that are spatially autocorrelated in the brain. Consistent with the intuition developed in Fig. 1, as the spatial autocorrelation increases, the false positive rates increase. Importantly, false positive rates are proportional to the average deviation in spatial autocorrelation between the original and rotated maps (*r* = 0.76). In summary, projecting brain maps to a spherical surface distorts distance relationships, yielding increased false positive rates.

### Targeted removal of poor nulls improves performance

If a specific spherical rotation (realization of the spin test null model) does not adequately preserve the spatial relationships in brain maps, then a straight-forward solution would be to remove it from the population of nulls. To quantify how well a rotation preserves distance relationships among vertices, we compute the correlation between vertex-to-vertex distance matrices of original and rotated surfaces (Fig. S1). Fig. S2 confirms that rotated maps with the poorest spatial distance alignment are those whose spatial autocorrelation deviates the most from the original map. We then repeat the experiments from the previous section, but gradually remove individual rotated maps that are poorly aligned with the original map.

Fig. 3a shows FPRs when applying this removal heuristic. As before, we quantify FPRs at varying levels of spatial autocorrelation. Different lines illustrate performance at different thresholds, from 0% removal to 90% removal. Note that, to ensure that all null populations are of equal size, we always sample 1 000 nulls out of the original 10 000-null ensemble (see *Methods*). As poorly-aligned spins are gradually removed, we observe lower FPRs. In these specific simulations (though the result may not generalize to all situations), we observe 5% FPRs at approximately 77.5% removal (Fig. 3b). At greater removal rates, FPRs dip below the desired 5% level; Fig. 3c shows that this might be because the remaining null realizations resemble the original map too closely and do not sufficiently sample the null space. In summary, removing poorly-aligned rotated maps potentially results in improved performance of spatial permutation tests but entails heuristic intervention in the sampling procedure that may be sub-optimal.

**Figure 3.**
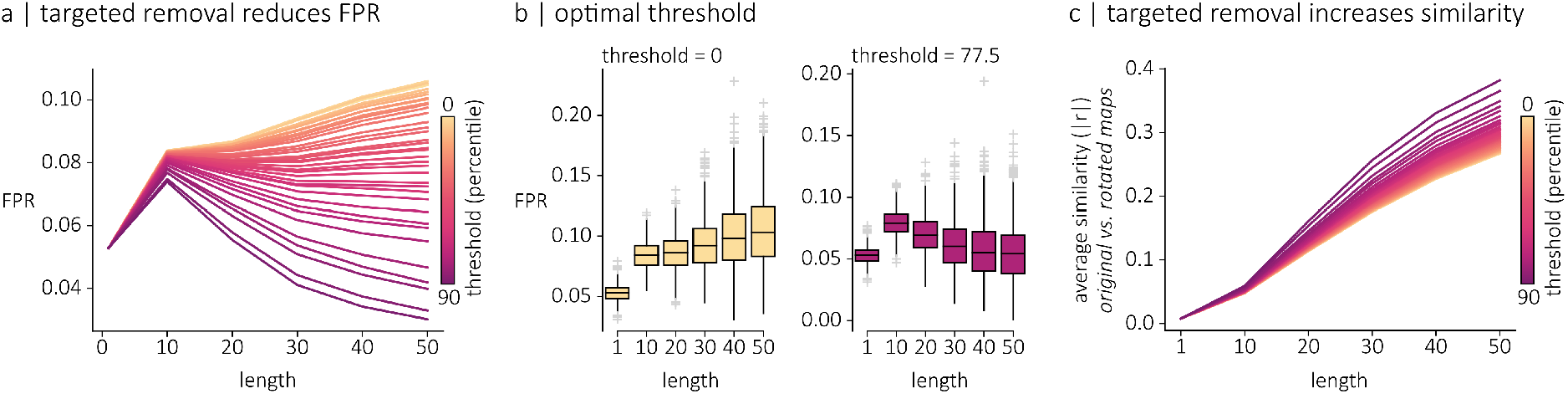
Targeted removal of poor nulls improves performance. (a) Statistical performance of the spin test (FPR) following the targeted removal of suboptimal rotations. Individual lines represent the average FPR across simulated maps, for each autocorrelation level. Line colors indicates the threshold (in percentile) for the removal of rotations, ranging from 0 to 90. (b) Distribution of false positive rates quantified for each map individually, across autocorrelation levels when no threshold is applied (right) and when a threshold of 77.5 is applied (left). (c) Average similarity (absolute Pearson coefficient) between original and permuted maps for each autocorrelation levels, with each line representing a different threshold for the removal of rotations.

## DISCUSSION

Spatial correlations between brain maps are a fundamental component of scientific discovery in brain imaging. Here we explored why a popular spatial null model for assessing statistical significance does not adequately control false positive rates. We then show how a straight-forward modification of the method can reduce false positive rates.

We find that false positive rates in spatial permutation tests can be traced back to the initial step of projecting a brain surface to a sphere. Inflating a brain surface to a sphere induces metric distortions due to cortical curvature [38]. Most inflation algorithms attempt to minimize distortions, but they are still present. As a result, patterns of autocorrelation on a brain surface are distorted when projected to a sphere. Importantly, we confirm that subsequent angular rotations of the spherical representation are not problematic, as they are isometric transformations.

This work is part of an emerging appreciation of the fact that the brain surface is irregular. Namely, the standard preprocessing step of mapping volumetric voxels to a highly folded surface yields uneven surface vertex spacing, such that neighbouring vertices are physically closer in sulci than in gyri. This uneven sampling can be suboptimal [39] and can introduce a “gyral bias” in fMRI analyses [40]. The distortions that we observe when mapping cortical surfaces to spheres and which lead to inflated p-values in the spin test are directly related to this uneven sampling. In this sense, our findings point to a need to develop methods that more accurately represent and manipulate cortical geometry. How to quantify features of local geometry on the brain surface, such as autocorrelation [41] or wave patterns [42], is an open question in the field.

We explore a simple heuristic to reduce the effect of ill-preserved spatial autocorrelation on false positive rates. We find that some realizations of spins are better than others, and trace the inflated false positive rates to specific realizations that poorly preserve spatial autocorrelation of the empirical map. We show that targeted removal of deviant individual null realizations can reduce false positive rates to desirable levels. Conceptually, this procedure can be thought of as performing posthoc quality control on the spatial null generating process. Ultimately, the goal of the spatial randomization procedure is to sample the wider null space of brain maps with similar autocorrelation [31]; the proposed “fix” helps to prune and narrow down this space such that it only includes realizations of sufficient quality, i.e. sufficiently similar autocorrelation to the empirical map.

Going forward, what spatial autocorrelation-preserving method should researchers use? On one hand, spin tests are fast, easy to apply, and the same “spin” can be applied uniformly to all maps in a given sample. On the other hand, the proposed solution for reducing false positives requires additional experimenter intervention and may potentially bias sampling of the null space. Fortunately, this is an active field of research and there exist numerous alternative methods to randomize brain maps while preserving spatial autocorrelation [16, 28, 30, 43]. Ultimately, the field is still in its infancy and it is too early to draw overly prescriptive conclusions. More research is warranted in this domain, not only on null models but on the geometric features of the brain more generally.

## METHODS

### Spatially autocorrelated maps

To evaluate the statistical performance of the spin test, we ran simulation experiments using spatially autocorrelated maps generated with the GSTools python toolbox [44]. The spatial properties of these maps were derived from a Gaussian variogram model defined as follows:

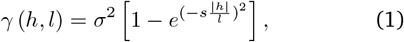

where *h* denotes the Euclidean distance between vertices. The parameter *s* is a rescaling factor, set to the toolbox’s default value of 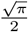, *l* denotes the length of the model and *σ*^2^ denotes the variance of the model. From this variogram model, a spatially autocorrelated random map *u*(**x**) can be generated using the following calculation, known as the randomization method [45, 46]:

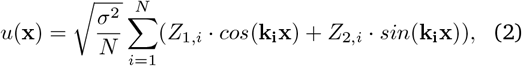

where *N* is the number of Fourier mode used. Here, *N* was set to 1 000. *Z*_*i*,*j*_ are random samples drawn from a normal distribution and **k**_**i**_ are samples from the spectral density distribution of the covariance model. The range of the variogram (i.e. distance at which the maximal variance is attained) approximately corresponds to 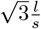. Thus, the larger the length parameter, the greater the spatial autocorrelation of the maps generated. To evaluate the performance of the spin test across levels of autocorrelation, 1 000 random fields were generated for six length values ranging from 1 to 50. To compare the performance of the spin test across surfaces, maps were generated on two different meshes of the left hemisphere’s fsaverage5 template [47], namely the pial and the spherical surface meshes.

### Moran’s I

The spatial autocorrelation of the maps considered in this work was quantified using Moran’s I [48, 49]:

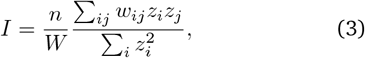

where *n* corresponds to the number of spatial units (vertices), *w*_*ij*_ corresponds to the spatial weight quantifying the spatial relationship between vertices *i* and *j, W* corresponds to the sum of all weights *w*_*ij*_ and 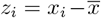. The spatial weights were defined as the inverse of the row-normalized Euclidean distance matrix between pairs of vertices.

### The spin procedure

The results presented in this work relied on the fsaverage5 cortical mesh from FreeSurfer [50]. Spatially autocorrelated maps generated on the pial surface of the brain were first projected to the spherical mesh. This step was not necessary for spatially autocorrelated maps generated directly on this spherical mesh. Then, random rotations were applied to the spherical mesh using QR decomposition. The vertices on the rotated mesh were then matched to vertices on the original spherical mesh using a nearest neighbor approach. This process was repeated until a total of 10 000 realization (spins) were generated. For all experiments, only a subset of 1 000 spins were used. The 10 000-null ensemble was initially sampled at random. Then, for the later experiments, a certain percentage, ranging from 0 to 90%, of the lowest-quality spins were removed from this null ensemble, and 1 000 spins were then sampled from this thresholded ensemble.

### Statistical performance of the spin test

The statistical performance of the spin test was evaluated using the spatially autocorrelated maps described in the first section of the *Methods*. For each spatial autocorrelation level (defined by the length parameter), a total of 1 000 maps were generated. The Pearson correlation between each pair of maps {**X, Y**} was computed, and its significance was evaluated using the spin test. Namely, a total of 1 000 random rotations were applied to the map **X**. Each rotated map was then correlated with the map **Y** to generate a null distribution of correlation coefficients. The *p*-value of the correlation coefficient between {**X, Y**} was then evaluated relative to this null distribution. For each map **X**, this procedure was repeated across all maps **Y**, resulting in a distribution of 999 *p*-values. The false positive rate of **X** was then computed as the ratio of significant relationships (*p <* 0.05 threshold).

### Quantifying the quality of spins

The spin procedure involves projecting a cortical surface (e.g. pial) *P* to a sphere *S*, rotating the sphere *S* to a sphere *S′*, then projecting the sphere *S′* back to a cortical surface *P′*. By concatenating all transformations, we get a mapping from a cortical surface *P* to a cortical surface *P′*. The quality of this mapping between the two cortical surfaces was quantified as the Pearson correlation between the original and transformed distance matrices (Fig. S1).

## DATA AND CODE AVAILABILITY

The data and code used to conduct the analyses and generate the figures presented in this paper is available at https://github.com/netneurolab/bazinet_spins and directly relies on the following open source Python packages: NumPy [51], Scipy [52], Matplotlib [53], GSTools [44], NiBabel [54], PyVista [55] and neuromaps [1].

## ACKNOWLEDGMENTS

We thank Filip Milisav, Justine Y. Hansen, Eric G. Ceballos, Yigu Zhou, Asa Farahani, Andrea Luppi and Moohebat Pourmajidian for their comments and suggestions on the manuscript. VB acknowledges support from the Natural Sciences and Engineering Research Council of Canada (NSERC), the Fonds de Recherche du Quebec - Nature et Technologie (FRQNT), the Healthy Brains for Healthy Lives initiative and Brain Canada. BM acknowledges support from the Natural Sciences and Engineering Research Council of Canada (NSERC), Canadian Institutes of Health Research (CIHR), Brain Canada Foundation Future Leaders Fund, the Canada Research Chairs Program, the Michael J. Fox Foundation, and the Healthy Brains for Healthy Lives initiative. The funders had no role in study design, data collection and analysis, decision to publish or preparation of the manuscript.

---

**Figure S1.**
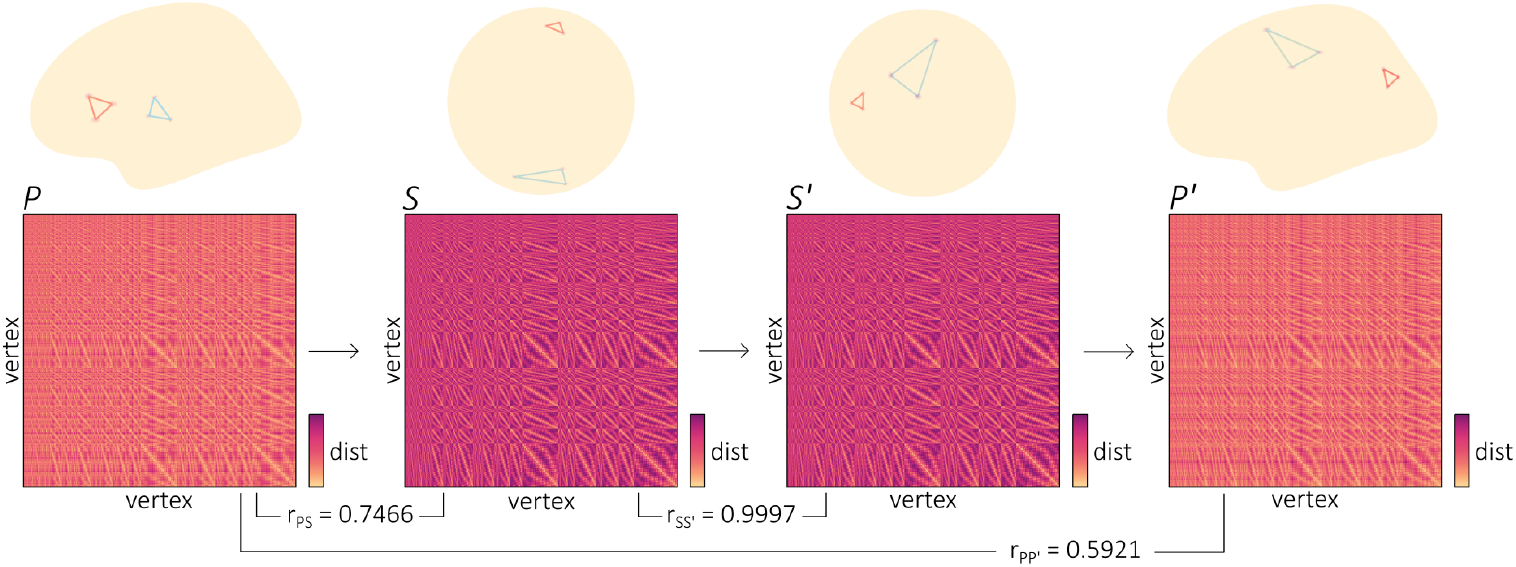
Quality of spins. The spin procedure involves projecting a cortical surface *P* to a sphere *S*, rotating the sphere *S* to a sphere *S′*, then projecting the sphere *S′* back to a cortical surface *P′*. The quality of each transformation can be quantified as the Pearson correlation between the distance matrix in the original space and the distance matrix in the transformed space. For the first transformation *P* → *S*, we have *r*_*PS*_ = 0.7466. The fact that *r*_*PS*_ is not equal to 1 indicates that the projection is not isometric and that distortions have been introduced. For the second transformation *S* → *S′*, we evaluated the quality of a random rotation obtained via QR decomposition and found: *r*_*SS*_*′* = 0.9997, which confirms that a rotation of the sphere is an isometry. Note that *r*_*SS*_*′* is not exactly equal to 1 because the spherical mesh is a recursively subdivided icosahedron, and not a perfect sphere. By concatenating each transformation, we ultimately obtain a transformation *P* → *P′*, which captures the entire spin procedure. For that specific spin instance, we have *r*_*PP*_ *′* = 0.5921.

**Figure S2.**
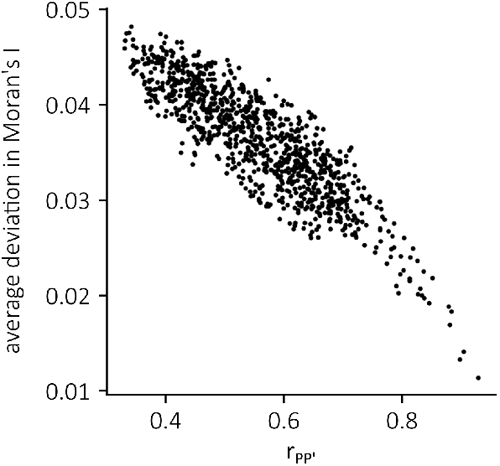
Rotations of poorer quality are associated with larger deviations in Moran’s I. The quality of each rotation was evaluated as the Pearson correlation between the distance matrix in the original space and the distance matrix in the transformed space (*r_P P_'*). The quality of each rotation was then compared to the average deviation in Moran’s I between original maps and maps permuted with that specific rotation. Namely, each rotation was applied to 1,000 spatially autocorrelated maps (length=50) and, for each map, the absolute difference between its Moran’s I following the rotation and its Moran’s I prior to the rotation was evaluated. For each rotation, its average deviation in Moran’s I was then computed as the mean of this absolute difference, across all maps. We find a strong negative relationship between the quality of a rotation and the average deviation in Moran’s I associated with that rotation (*r* = −0.88).

